# A portable 3D-printed near-infrared spectrometer to screen maize for deoxynivalenol and zearalenone contamination

**DOI:** 10.1101/2025.10.23.684207

**Authors:** Mariane Zabotto Evangelista, Dedy L. Nadeak, Rudolf Krska, José Tadeu Aldrigue, Bernardette Cajaiba Oliveira Rossiti, José Alexandre Melo Demattê, Borges Marfrann Dias Melo, Stephan Freitag, Marcos Livio Panhoza Tse

**Author notes:** Corresponding authors (SF), (MLPT). Current Address: Department of Agricultural Sciences, Institute of Bioanalytics and Agro-Metabolomics (BOKU – IFA), Tulln, Austria. Current Address: Department of Animal Production and Preventive Veterinary Medicine, São Paulo State University “Júlio de Mesquita Filho”, School of Veterinary Medicine and Animal Sciences (UNESP – FMVZ), Botucatu, Brazil.

## Abstract

Mycotoxins are toxic metabolites found in grains and cereals, posing a severe potential risk to human and animal health and significantly disrupting animal production, particularly in industries like pig farming. Current laboratory methods for mycotoxin analysis are expensive and time-intensive, creating a need for faster, more accessible solutions. This study set out to address this challenge by developing a portable spectrometer prototype based on the near-infrared (NIR) region and designing a chemometric model to classify ground maize as either non-compliant (NC) or compliant (C) for zearalenone (ZON) and deoxynivalenol (DON). A total of 259 naturally contaminated maize samples, collected over two years from diverse European regions, were analyzed using both liquid chromatography-tandem mass spectrometry (LC-MS/MS) and the prototype device. Spectral data were preprocessed using the first derivative, and a partial least squares discriminant analysis (PLS-DA) model was developed to classify ZON and DON levels into NC and C categories. Thresholds of 100 µg/kg for ZON and 500 µg/kg for DON were used to define compliance. The PLS-DA model showed good performance for ZON, achieving a classification accuracy of 86.3%. However, for DON classification a rather limited accuracy of 66.75% was achieved. While the DON model could identify NC samples it struggled with C samples. Despite these challenges, the results highlight the potential of a portable NIR spectrometer, combined with straightforward preprocessing and PLS-DA modeling, as a rapid, cost-effective screening tool for detecting mycotoxins in maize. The presented approach could significantly simplify mycotoxin monitoring, offering a practical solution to safeguard public health and enhance agricultural productivity.

## Introduction

Mycotoxins are secondary metabolites of filamentous fungi found in food and feed [1]. In addition to their toxic effect on humans, the occurrence of the mycotoxins zearalenone (ZON) and deoxynivalenol (DON) in feed causes negative effects on animal production, mainly in monogastric animals, such as poor feed absorption, liver damage, reduced growth, changes in the immune system, and reproductive problems [2]. There is a wide variety of methods available for mycotoxin analysis, but most of them come with the disadvantages of complex sample preparation, the use of a laboratory environment, expensive equipment, and the need for a specialist to perform the analysis [3, 4, 5, 6].

Consequently, small companies and farmers have difficulty accessing these laborious, time-consuming, and expensive laboratory tests. However, rapid analysis is often required for decision-making, which fuels the need for rapid on-site mycotoxin screening [7].

Rapid and easy-to-use screening approaches based on infrared spectroscopy can fill this gap [8, 9]. Near-infrared (NIR) spectroscopy has long been used for food and feed analysis [10], and a wide variety of portable NIR spectrometers are available [11].

Recent studies have shown the potential of portable spectrometers as an alternative method to screen mycotoxins on a semi-quantitative basis. Portable NIR spectrometers paired with chemometric tools show also some promise for rapid screening of mycotoxins in grains and cereals, such as peanuts and maize [12, 13].

In addition, 3D printing has found its way into the analytical laboratory [14, 15]. 3D-printing has been used to build portable spectrometers or accessories for benchtop devices [16, 17, 18] a sample preparation kit for mycotoxin screening, among many other applications [19] [7].

This study aimed to build a reliable portable and cost-efficient 3D-printed portable NIR spectrometer prototype to screen maize for ZON and DON. We developed a method validation of this screening tool by combining NIR spectra, liquid chromatography tandem mass spectrometry (LC-MS/MS) data, and chemometric modeling to classify maize into non-compliant (NC) and compliant (C) samples at a threshold of 100 µg/kg for ZON and 500 µg/kg for DON, based on Europe legislation of Commission Regulation (EC) n° 1881/2006.

## Materials and Methods

### Development of the portable near-infrared spectrometer

The technical drawings of the spectrometer, consisting of several parts, were designed using Inventor® Professional 2025 (Autodesk®) to achieve a compact device.

The housing of the spectrometer and the sample presentation prototype was manufactured by 3D printing using an Original Prusa I3 MK3 printer (Prusa Research by Josef Prusa) using the PrusaSlicer slicing software, version 2.7.4, 64 bits (Prusa Research by Josef Prusa; Slic3r open-source project), in 1.75-mm-diameter polylactic acid filament. During the slicing of the 3D models, auxiliary supports were created in all cavities and curves with angles from 30° were created. The filament flow during printing was set to the extrusion multiplication factor of 1.03, the height of the print layers was 0.15 mm, and the nozzle temperatures and the plate temperature were 210°C and 60°C, respectively.

The prototype, measuring 14.5 x 15 x 14 cm, was assembled by joining the printed parts using fastener sets. The spectrometer module has two connections: one USB output for transmitting the collected data and a 5V 2A power connector. The NIR spectrometer module was mounted to the top of the sample handling interface so that the measuring interface was 8 mm above the sample, as shown in Figure 1.

**Fig 1.**
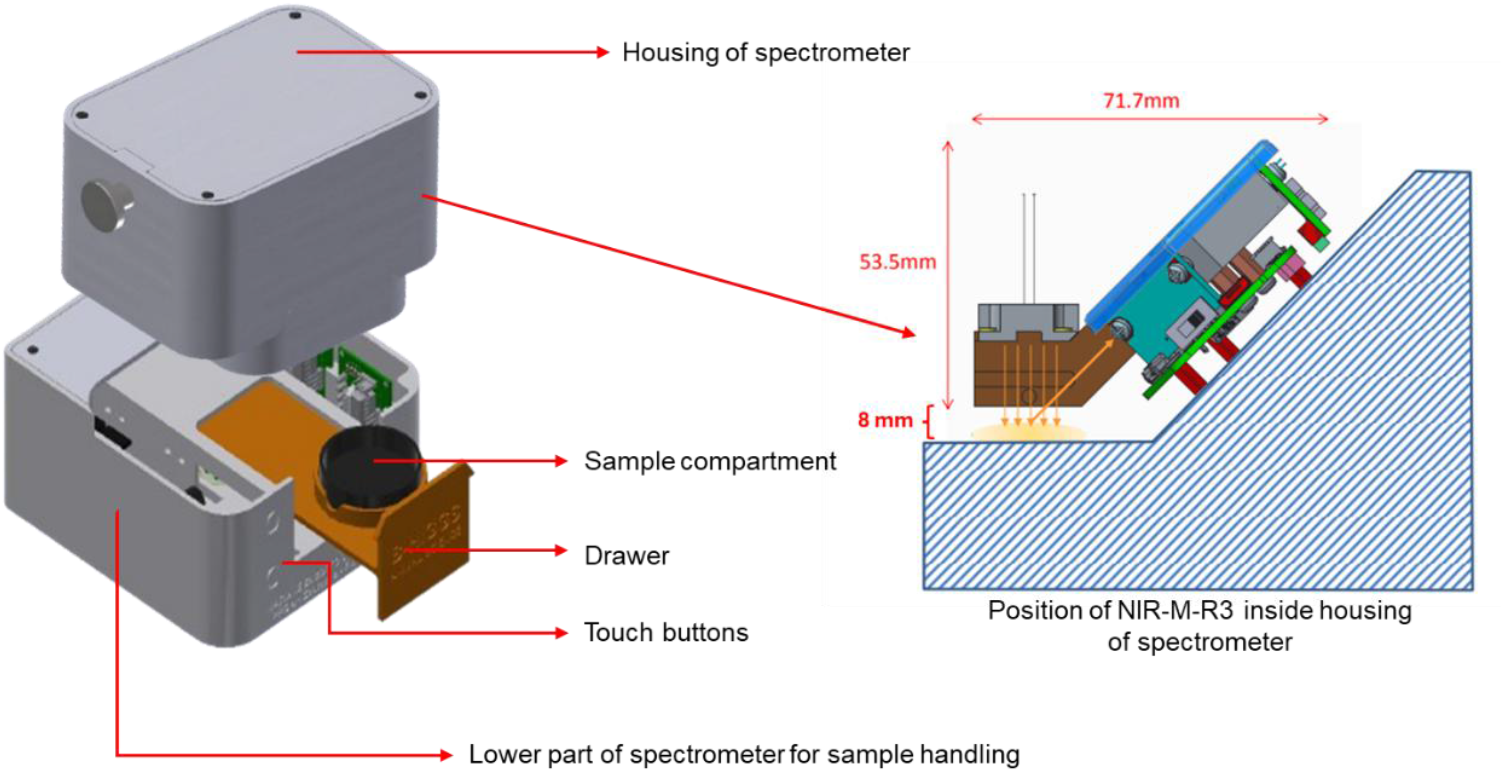
Schematic drawing of the 3D-printed portable prototype and its functional parts with the NIR module position configuration. Source: Adapted from [22].

The research and development spectrometer model was the NIR-M-R3 from InnoSpectra, with DLP (Digital Light Processing) technology, measuring 75.5 mm x 53.5 mm x 71.7 mm, operating in diffuse reflection mode with a spectral coverage ranging from 900 to 1700 nm, wich has been used in other studies [20, 21].

The assembly and fixing of all components (Table 1) for the prototype were performed manually.

**Table 1.** Components used in the assembly of the portable prototype.

| Amount | Component | Technical description |
| --- | --- | --- |
| 01 | NIR spectrometer module | Model NIR-M-R3, 900-1700 nm, diffuse reflection, TI DLP2010NIRAFQJ 0.2 WVGA near-infrared DMD, detector 1 mm InGaAs, tungsten lamp 5W. |
| 02 | Stepper motor | Model 28BYJ-48 + ULN2003 controlator |
| 02 | Microswitch | Limit switch with 01 normal open contact, 01 normal closed contact, and 15 A 125/250 VAC. |
| 01 | Arduino board | Development board AVR Arduino Uno Rev 3 |
| 01 | Buzzer | Piezo sounder for Arduino, 2 GND, 1 VDC external, supply voltage: 3,3-5V. |
| 02 | Touch button | Capacitive touch sensor for Arduino, operating voltage: 2,0-5,5V. |
| 20 | Wire jumper | 20 cm interconnect cables for male/female prototyping 20 cm. |
| 10 | Fastener set | Threaded metal inserts, screws, and nuts |

The spectrometer module is equipped with a tungsten lamp, a DMD wavelength selector, and a single-pixel InGaAs detector. The spectral range from 900 to 1,700 nm covers overtones of C-H, O-H and N-H molecular vibrations [23].

The DMD captures a single wavelength and reflects the desired wavelengths to the detector, providing greater precision to the physical signal readings [24]. The capture of one wavelength at a time is due to the use of a Hadamard standard correction code, which detects and corrects errors during the transmission of the spectra. Each pattern simultaneously turns on 50% of the micromirror array pixel, directing a much larger signal to the detector.

Two stepper motors were used for sample handling and presentation, i.e., to rotate the samples during measurements and to position the samples underneath the NIR source and detector in a reproducible manner. This automated sample presentation approach was controlled by an Arduino board linked to a buzzer and two microswitches. The total cost of producing the prototype was USD 2,800.00.

### Maize Sample Set and Liquid Chromatography Tandem Mass Spectrometry

The samples and the corresponding LC-MS/MS reference values were supplied by the Agrana Research and Innovation Center (ARIC) in Tulln and consisted of 259 ground maize samples of approximately 300g each from different European countries with the corresponding mycotoxin values.

All samples were analyzed using liquid LC-MS/MS according to the method of Varga et al. [25] by the ARIC to determine the concentration of the mycotoxins ZON and DON. All samples were milled using a Thermomix (Vorwerk, Wuppertal, Germany), and 10 g was used for LC-MS/MS analysis. The ARIC reported a relative intermediate precision of 10% for DON and 15% for ZON.

### Acquisition of near-infrared spectra

NIR spectra from the maize samples were collected in the spectral range of 900–1700 nm using the developed prototype spectrometer described above.

The circular movement of the sample compartment provided by stepper motors was used to increase the representativity during spectral acquisition. The sample compartment was fixed on a moving stage, which allowed free and easy handling of the samples when opened.

The drawer was operated using touch-sensitive buttons connected to an Arduino microcontroller that manages the control signals. A dedicated firmware was developed to regulate the speed of the motors and the distance traveled by the sample compartment, programed to rotate 72° with each press of the corresponding touch button. Furthermore, the touch buttons were set to control the drawer’s opening and closing by simply pressing them.

The samples were placed into the sample compartment and spectra were collected using the NIR module software provided by (insert name of company): ISC WinForms SDK GUI, version 3.9.9.

The pre-recorded reference spectra provided with the spectrometer were used as the background, and the lamp stabilization time during spectra acquisition was set to 625 ms (standard), with 32 scans per measurement. The time required for spectra acquisition was approximately 20 s, and the Hadamard correction was used during the collection.

Each maize sample was subdivided into five subsamples of approximately 15g each and subjected to spectra collection with five repetitions (five rotations of the sample), totaling 25 spectra for each sample and 6.475 spectra in total.

### Data analysis

Data analysis and chemometrics were performed in R using RStudio, version 4.3.2 (Posit and R Foundation for Statistical Computing) and the R packages prospectr, ggplot2, ggpubr, and mdatools [26, 27, 28, 29].

In the first step, the 25 NIR spectra per sample were averaged, reducing the 6.475 to 259 spectra. This was done to ensure the representativity of the obtained NIR spectra.

A first-order gap derivative with a width of 3 and a segment size of 5 was used for spectral processing. PCA was used for exploratory data analysis. The dataset was split into training and test sets based on the Kennards-Stone algorithm, with 80% of the samples used to train the PLS-DS models and 20 % for validation. For ZON, a threshold of 100 µg/kg was used, and for DON, a threshold of 500 µg/kg was chosen to obtain relatively equally balanced classes. PLS-DA was used to establish models for the discrimination of non-compliant (NC) and compliant (C) samples. For ZON and DON, sperate models were created. The latent variables of the model PLS-DA parameters were selected using 10-fold cross-validation using a Venetian split of the training data. The two models’ performance was evaluated based on their ability to classify samples into NC and C, as follows:

- TNC: Samples found to be NC by LC-MS/MS and correctly predicted by the PLS-DA model as NC.
- TC: Samples found to be C by LC-MS/MS and correctly predicted as C by the PLS-DA model.
- FNC: Samples found to be C by LC-MS/MS and incorrectly predicted as NC by the PLS-DA model.
- FC: Samples found to be NC by LC-MS/MS and incorrectly predicted by the PLS-DA model as C.

The model’s ability to discriminate between NC and C was assessed using the following formulas:

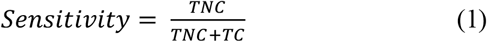

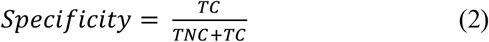

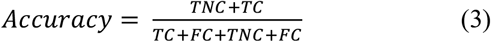

## Results and Discussion

### Spectra analysis

Figure 2 shows the averaged raw and the first derivative of the NIR spectra of the 259 different ground maize samples.

**Fig 2.**
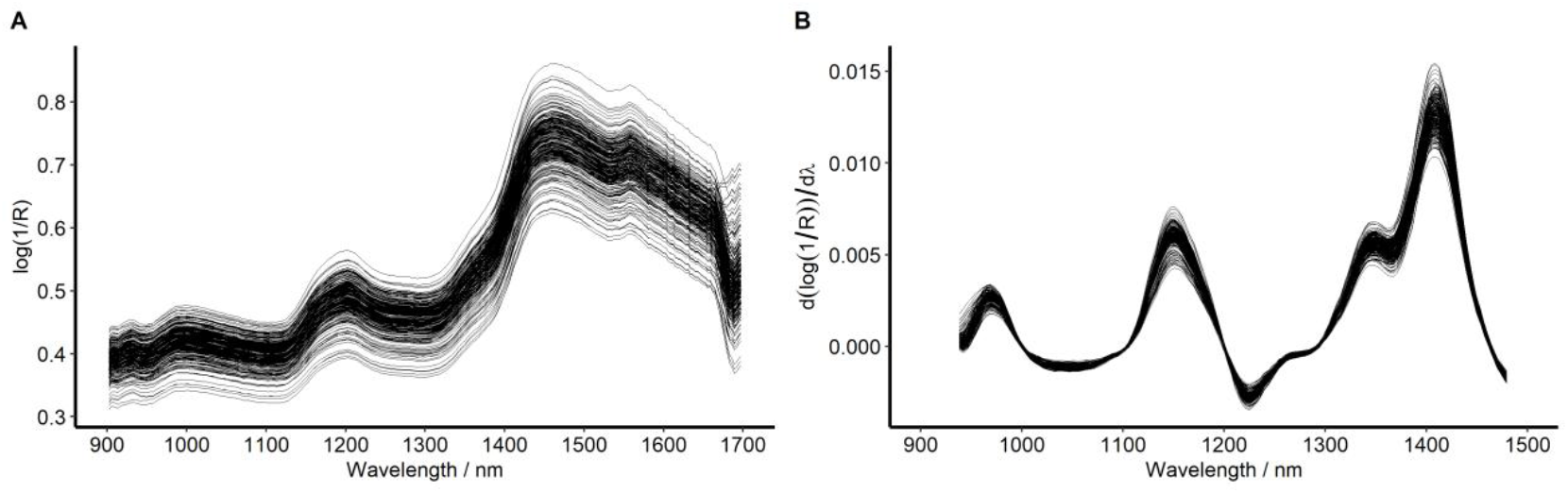
NIR Spectra. (A) Unprocessed spectra. (B) First derivative of the raw spectra after spectra truncation.

In Figure 2A, the spectra show variations in the baseline and slope, which can be attributed to the light scattering effects resulting from the differences in particle size or sample presentation [30]. These physical effects cause baseline shifts and amplitude variations between the spectra. Variations in particle size may explain the observed differences in shift and slope in the raw spectra [31, 32] and may have also affected the identification of properties and chemical changes in corn due to masking or highlighting bands [32].

Small variations are observed from 900 to 1100 nm, which can be attributed to aliphatic bonds in lipids and starch (C-H) and residual moisture (O-H harmonic combinations). The moderate peak in the 1150-1250 nm region corresponds to the first C-H overtones (CH_2_, CH_3_) of carbohydrates and lipids, which are associated with starch (amylose/amylopectin). The 1350-1450 nm region reflects the variation in moisture among samples, with strong water absorption (first overtone of the O-H stretching vibration). The 1500-1600 nm region shows the presence of proteins (N-H, mainly zein in corn) and polysaccharides (C-H) and may also indicate variations in crude protein content.

As shown in Figure 2B, spectral processing using the first derivative was applied in combination with spectral trimming to the 900-1500 nm range to reduce noise, particularly above 1500 nm in this device. After processing, a better resolution of the bands at 970–1000 nm, 1200 nm, and 1450 nm was observed, as well as a reduction in the offset seen in Figure 2A, indicating that pretreatment effectively eliminated the background interference and enhanced the spectral information, which was consistent with the published results of fumonisin B1 and B2 analysis in ground corn using a portable device [13].

According to the PCA presented in Figure 3, the first two components explain 88.54% of the total variance found in the processed NIR spectra (PC1: 66.76%; PC2: 21.78%). No separation was observed between the 2022 and 2023 harvest years or between the training and test sets. This indicates that the dominant spectral variability is linked to the variability of the sample set used and not to other effects. The two-year overlap should promote the development of robust chemometric models [10].

**Fig 3.**
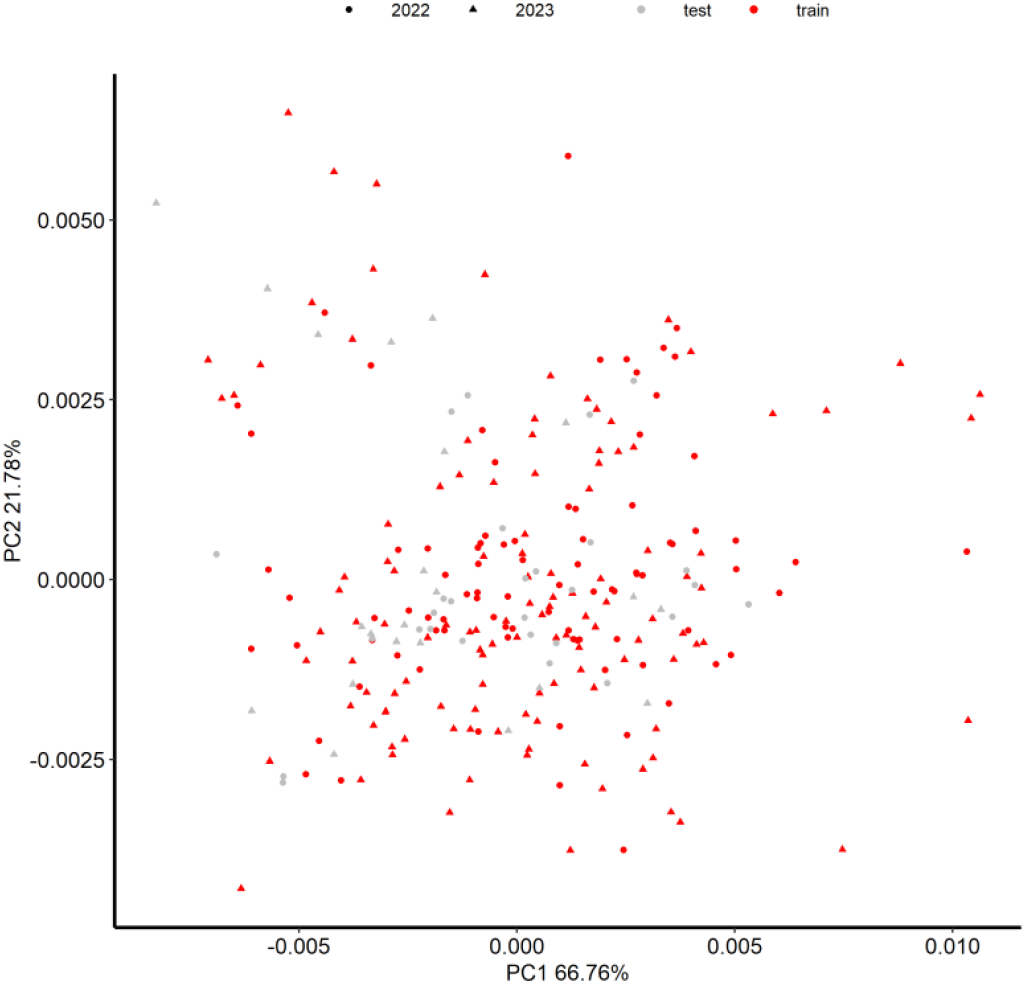
Principal component analysis of the first derivative spectra. Red and gray indicate the samples used as training data and testing the models, respectively.

The data were split into 80% and 20% for the training and validation sets, respectively, using the Kennard-Stone algorithm, as this method leads to a robust representation of the training set in the test set [33]. This pattern of overlap between the calibration and validation sets is consistent with the results reported in previous studies that also obtained representative data using similar set separation strategies in grains and other organic matter [34, 35, 36].

### PLS-DA for ZON and DON screening

Figure 4 shows the evaluation of the optimal number of latent variables (LV) aimed at balancing low error and high robustness in the resulting PLS-DA model. This analysis is shown separately for DON (Figure 4A) and ZON (Figure 4B). The red line represents the performance of the PLS-DA model during calibration (CAL) on the training set, while the blue line shows the model’s performance during CV using the training set.

**Fig 4.**
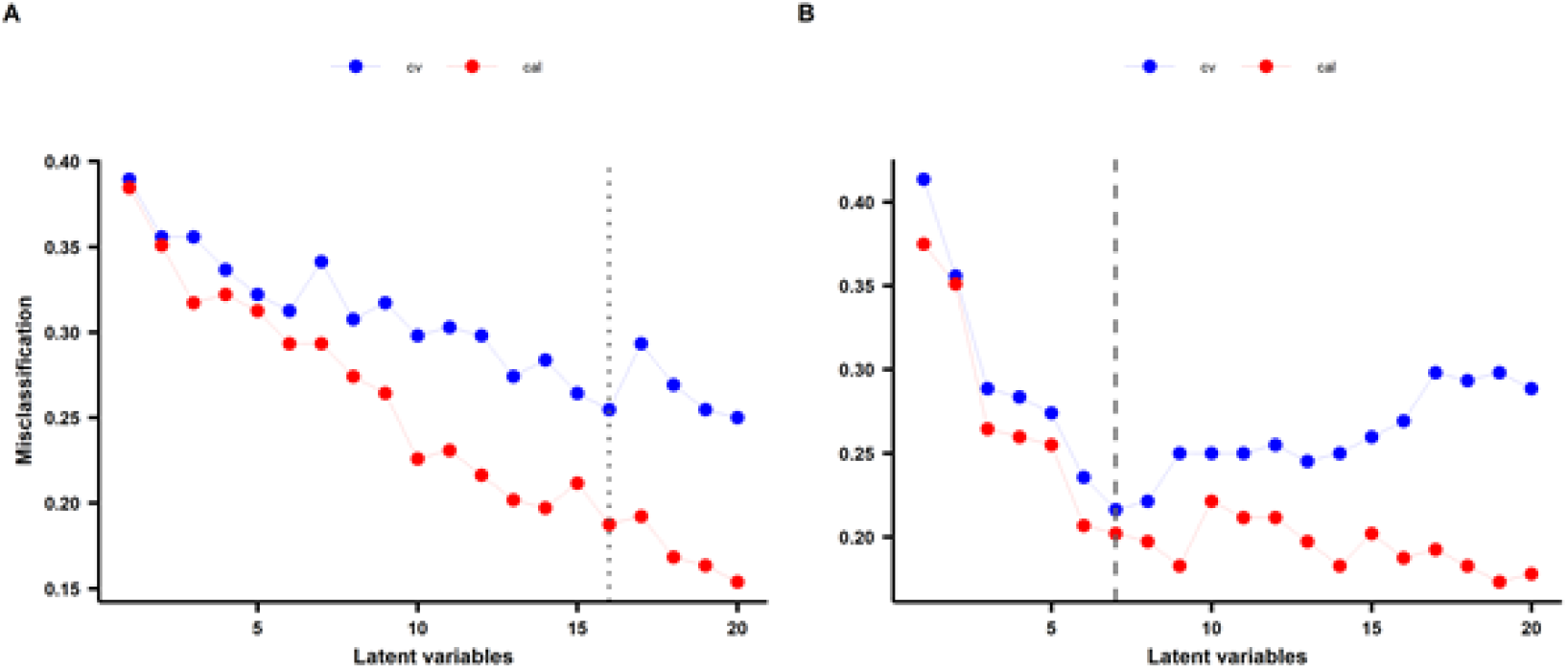
Plots of the misclassification rate. (A) DON. (B) ZON. cal = calibration, cv = cross-validation.

Figure 4 shows that the model’s performance during CV is less optimistic compared to simple calibration. It is well established that CV is used to tune models in a way that minimizes the prediction error while aiming to avoid overfitting the model to the training data [33, 37]. The models exhibited greater complexity for predicting DON contamination (LV = 16) than ZON (LV = 7). For DON, the misclassification rate decreases gradually with the number of LVs, whereas for ZON, a steeper decline is observed (Figure 4). Furthermore, the difference between the misclassification rates for calibration (CAL) and cross-validation (CV) is notably smaller in the ZON model than in the DON model. This suggests that PLS-DA is more effective at extracting information related to ZON contamination from the NIR spectra than DON contamination. A closer match between CAL and CV misclassification rates could indicate a better balance between model fit and generalization [38, 39]. This is underlined by the performance of the models on the validation samples (Table 2). The PLS-DA model for DON showed a lower accuracy of 66.7% when predicting the validation set, indicating the difficulty in maintaining performance outside the training set. A decrease in performance, i.e., an increase in misclassification, was observed, especially for compliant samples (specificity = 51.9%, FC = 14), indicating a low capability for recognizing compliant (low contaminated) samples. However, the sensitivity remained relatively high (83.3%), indicating that the model performs better in detecting NC samples. A similar performance was recently reported when a mid-IR approach was used [40]. To screen maize for DON, such a model might still be applicable, as NC samples should undergo confirmatory analysis [41].

**Table 2.** Results of the PLS-DA model for NC and C classes in ground maize for ZON (LV = 7) and DON (LV = 16).

|  |  | TNC | FNC | TC | FC | Specificity | Sensitivity | Accuracy |
| --- | --- | --- | --- | --- | --- | --- | --- | --- |
| **ZON** | *CV* | 60 | 21 | 103 | 24 | 0.831 | 0.714 | 0.784 |
|  | *V* | 22 | 3 | 23 | 4 | 0.88 | 0.846 | 0.863 |
| **DON** | *CV* | 80 | 30 | 75 | 23 | 0.714 | 0.777 | 0.745 |
|  | *V* | 20 | 13 | 13 | 14 | 0.519 | 0.833 | 0.667 |

TNC: true non-compliant; FNC: false non-compliant; TC: true compliant; FC: false compliant; CV: cross-validation; V: validation.

Similar results have been reported in previous studies, which confirm the difficulty of detecting DON due to its heterogeneous distribution and low concentrations in the matrix [42, 43], and suggest the use of variable selection methods and nonlinear models, such as random forest and convolutional neural networks, as a way to improve sensitivity for DON and model complexity [42, 44].

The accuracy obtained in this research was lower than those found in previous studies on DON discrimination in wheat grains using techniques combined with NIR (Vis/NIR, Vis/NIR/SWIR), which achieved an accuracy of 98% and 96%, respectively [45, 46]. These studies used naturally contaminated samples. Thus, as wheat is a matrix that commonly presents higher and more easily detectable DON contamination levels than other cereals even though maize can also be affected [47], our results suggest that the lower accuracy is mainly linked to the complexity of the natural contaminated maize matrix and to the lower levels of contamination observed. Moreover, the use of a portable prototype with a restricted spectral range compared to laboratory-grade instruments imposes additional discrimination challenges. Thus, models developed for DON screening using Savitzky–Golay + SNV preprocessing and PLSR/PLS-DA tend to perform better in wheat, with an accuracy of approximately 85% [48]. Other authors [49] have reported good results when analyzing NC and C samples for DON in maize using a threshold similar to that of this research (> or < 450 μg/kg, according to Brazilian legislation) in Brazilian wheat through dispersive NIR and FT-NIR, obtaining a correct classification rate between 85% and 87.5% for the classes. In contrast, the portable prototype developed in our study was used to collect spectra, which offers the advantage of being applicable directly in the field (in loco), became rapid and practical screening under real production conditions. Some studies have shown better DON detection results in barley. For instance, a study conducted to distinguish NC and C in barley samples with DON contamination (threshold 1250 μg/kg, according to European legislation) [50] achieved 90.9% specificity and 81.9% sensitivity.

The PLS-DA model obtained in this study showed better performance for classifying ZON-contaminated samples (accuracy = 86.3%), with a better ability to recognize both C and NC samples (specificity = 88%; sensitivity = 84.6%), compared with the DON model. High accuracy (100%) was also reported in a study in which the PLS-DA model developed with second derivative preprocessing was applied to the analysis of hulled barley [51]. The results obtained in this study are consistent with those of previous studies employing similar techniques, including NIR-HSI and PLS-DA, for ZON screening in cereals (accuracy above 85%) [43, 52]. This might be explained by the stronger correlation between the NIR spectra and ZON contamination, which might cause more pronounced alterations in the matrix [43, 53].

The results obtained are similar to those of a study that used a portable spectrometer with 900-1700 spectral range and PLS-DA modeling to screen fumonisin B1 and B2 in maize and obtained a high accuracy of 88% (85% sensitivity; 91% specificity) [13]. Their results indicate that a portable NIR spectrometer coupled with chemometrics had great participation in that, being a potential strategy for monitoring FB contamination in maize. A study that used a portable NIR device to screen aflatoxin B1 in peanuts, established after Beluga whale optimization and iteratively variable subset optimization (BWO-IVSO) algorithms [12], reported good model prediction and generalization performance. The results showed that the use of portable spectrometers associated with appropriate chemometric methods has a favorable application prospect in the quantitative analysis of mycotoxins.

Figure 5 summarizes the performance of the model by classifying the validation samples of the test set as compliant (C) or non-compliant (NC). Figures 5A and 5C show the classes found by LC-MS and the classes predicted by the model for ZON and DON, respectively. Figures 5B and 5D display the concentrations obtained by LC-MS/MS of the misclassified samples, with the red line indicating the threshold and the dotted line indicating the 95% confidence interval of the threshold of the LC-MS method. These boundaries are based on the expanded uncertainty (*U*) using a coverage factor (*k*) of 2 and calculated in accordance with the Eurachem/CITAC guide as follows [54].

**Fig 5.**
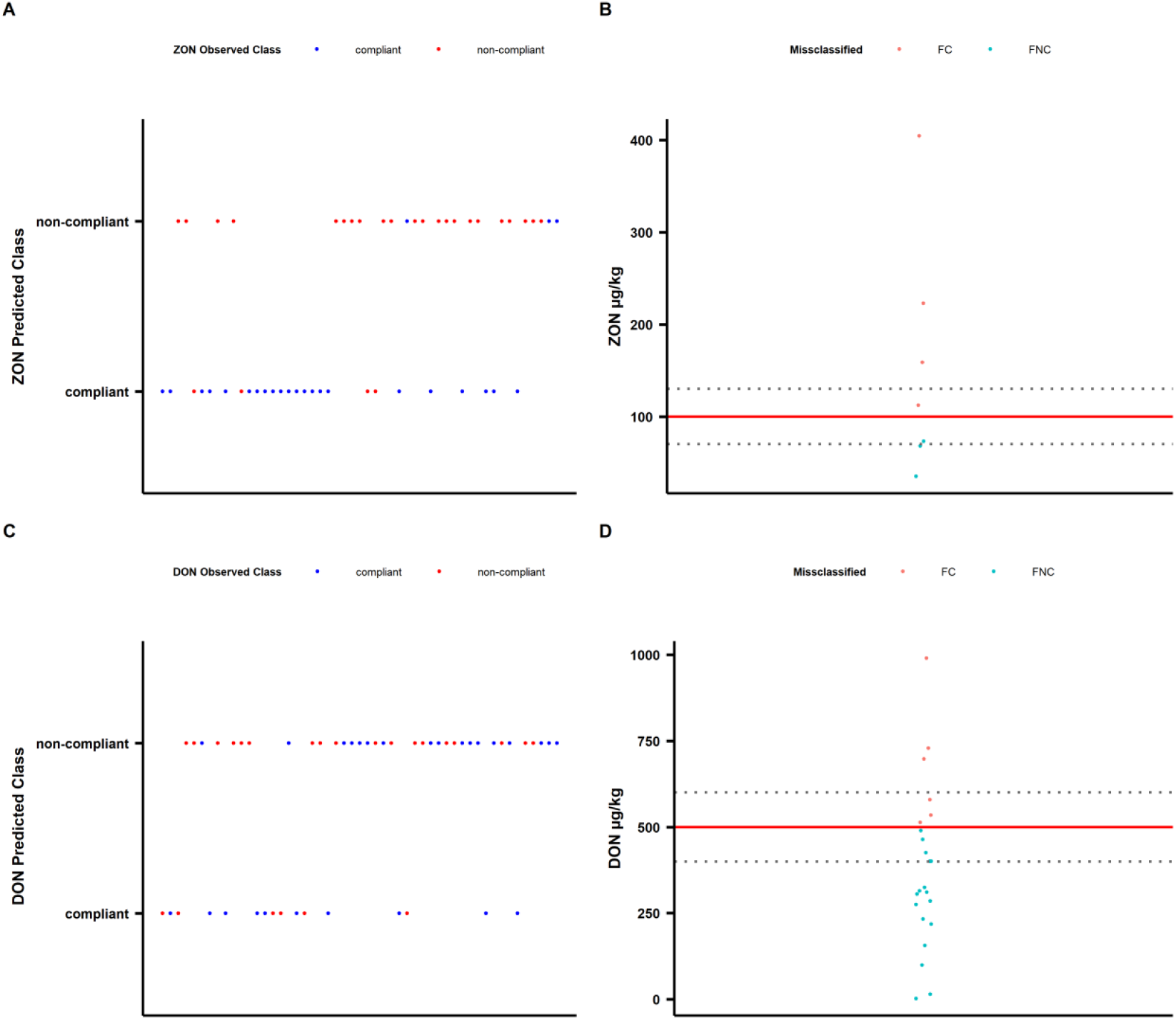
Performance of the PLS-DA model in discriminating maize samples. (C) Compliant. (NC) noncompliant. Red line = validation set with the discrimination thresholds, dashed gray lines = the confidence interval of the classification threshold, FC= false compliant, FNC = false non-compliant.

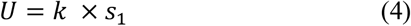

Where *U* is the expanded uncertainty of the measurement result, *k* is the coverage factor associated with the desired confidence level (typically k = 2 for approximately 95% confidence); *s1* is the combined relative uncertainty associated with the analyte.

Based on the provided relative intermediate precision (*S*_*I*_), with a confidence interval of 95% a lower limit of 70 µg/kg and an upper limit of 130 µg/kg was approximated for ZON. For DON, a lower limit of the confidence interval of 400 µg/kg and a higher limit of 600 µg/kg were assumed.

Figure 5 shows that the ZON model performs better. This may be due to matrix changes associated with ZON contamination generating more consistent and detectable spectral signatures compared to DON.

A large portion of the misclassified samples (both FC and FNC) are concentrated near the threshold (Figures 5B and D), which is expected because the reference value also has a certain error. Studies using HSI and pixel-based analysis have shown that these errors are typically concentrated in borderline zones, reinforcing the need for caution when interpreting screenings near the limit [48, 44]. In general, a relative standard deviation of <25% for ZON and <20% for DON would be still accepted by the EC in the concentration range of the thresholds [55]. Although the reported errors for the LC-MS/MS method are not that high, not all of the misclassified samples are within the CI of the threshold. This might be linked to the fact that the error introduced by sampling mismatches is not included, i.e., the sample used for NIR is not representative of the LC-MS/MS portion. However, sampling is the biggest source of error in mycotoxin analysis [56]. Furthermore, the presence of atoxigenic Fusarium strains might also lead to sample misclassification.

In summary, the applied methodology (first derivative preprocessing followed by PLS-DA) seems to be suitable for effective ZON screening but showed limitations for DON. Therefore, to increase robustness in DON screening, as recommended in the literature, sample representativeness (harvests, regions, natural contamination) should be increased, variable selection could be applied, and nonlinear algorithms should be tested along with external validation to ensure true generalization [48, 44, 42, 43].

## Conclusions

We demonstrated that a portable 3D-printed prototype NIR spectrometer can be used to screen maize for DON and ZON contamination in an easy and cost-effective way. By employing a simple spectra preprocessing strategy combined with PLS-DA modeling, we found that ZON screening is more straightforward than DON screening. In addition, we emphasize the importance of understanding the error associated with the reference values used. In future studies, we aim to enhance the prototype’s robustness, expand the sample set, and further improve the mycotoxin screening model’s performance. Our study highlights that mycotoxin detection using a custom-built portable NIR spectrometer appears feasible, provided that a robust screening calibration is developed to effectively manage the diversity of samples encountered in real-world scenarios.

## Acknowledgements

The authors would like to thank the School of Veterinary Medicine and Animal Sciences (FMVZ-UNESP), the University of “Luiz de Queiroz” College of Agriculture (ESALQ-USP), and the Institute of Bioanalytics and Agro-Metabolomics (BOKU – IFA Tulln, Austria) for their support during the development of this research. The authors are also grateful to Thomas Karner (ARIC) for supplying the maize samples and corresponding mycotoxin reference values.

## References

1. Gruber-Dorninger C, Jenkins T, Schatzmayr G. Global Mycotoxin Occurrence in Feed: A Ten-Year Survey. Toxins. 2019; 11, 7, 1–25. doi: 10.3390/toxins11070375.

2. Bunzen S, Haese D. Controle de micotoxinas na alimentação de aves e suínos. Rev Eletr Nutritime. 2006; 3, 1, 299–304.

3. Singh J, Mehta A. Rapid and sensitive detection of mycotoxins by advanced and emerging analytical methods: a review. Food Sci Nutr. 2020; 8. doi: 10.1002/fsn3.1474.

4. Yang J, Li J, Jiang Y, Duan X, Qu H, Yang B. Natural occurrence, analysis, and prevention of mycotoxins in fruits and their processed products. Crit Rev Food Sci Nutr. 2014; 54, 1, 64– 83. doi: 10.1080/10408398.2011.569860.

5. Pascale M, Visconti A. Overview of detection methods for mycotoxins. In: Leslie JF, Bandyopadhhyay R; Visconti A, organizers. Mycotoxins: detection methods, management, public health and agricultural trade. 1 ed. Wallingford: CABI; 2008. pp. 171–181.

6. Thway R, Salimi-Moosavi H. Evaluating the impact of matrix effects on biomarker assay sensitivity. Bioanalysis. 2014; 6, 8, 1081–1091. doi: 10.4155/bio.14.55.

7. Csenki E, Mikulás V, Freitag S, Fomina P, Hlavatsch M, Femenias A, et al. Stakeholder assessment for mycotoxin analysis: exploring the demand along the European food supply chain. World Mycotoxin J. 2023; 16, 287–299. doi: 10.1163/18750796-20232883.

8. Freitag S, Sulyok M, Logan N, Elliott CT, Krska R. The potential and applicability of infrared spectroscopic methods for the rapid screening and routine analysis of mycotoxins in food crops. Compr. Rev. Food Sci. Food Saf. 2022; 21, 6, 5199–5224. doi: 10.1111/1541-4337.13054.

9. Freitag S (2025). Infrared spectroscopy for mycotoxin screening: Advantages and limitations. In: Mycotoxins (pp. 249–266). doi: 10.1163/9789004724969_012.

10. Williams P, Antoniszyn J, Manley M. Near-infrared Technology: Getting the best out of light. 1. ed. African Sun Media, SUN PReSS; 2019. 311 p.

11. Crocombe RA. Portable Spectroscopy. Appl. Spectrosc. 2018; 72, 12, 1701–1751. doi: 10.1177/0003702818809719.

12. Li J, Deng J, Bai X, Monteiro DGN, Jiang H. Quantitative analysis of aflatoxin B1 of peanut by optimized support vector machine models based on near-infrared spectral features. Spectrochim. Acta A Mol. Biomol. Spectrosc. 2023; 303, 123208. doi: 10.1016/j.saa.2023.123208.

13. Shen G, Kanga X, Sua J, Qiua J, Liua X, Xua J, et al. Rapid detection of fumonisin B1 and B2 in ground corn samples using smartphone-controlled portable near-infrared spectrometry and chemometrics. Food Chem. 2022; 384, 132487. doi: 10.1016/j.foodchem.2022.132487

14. Pamidi AS, Spano MB, Weiss, GA. A Practical Guide to 3D Printing for Chemistry and Biology Laboratories. Curr protoc. 2024. doi: 10.1002/cpz1.70036.

15. Carrasco-Correa EJ, Simó-Alfonso EF, Herrero-Martínez JM, Miró M. The emerging role of 3D printing in the fabrication of detection systems. TrAC, Trends Anal. Chem. 2021; 136, 116177. doi: 10.1016/j.trac.2020.116177.

16. Baumgartner, B, Freitag S, Lendl B. 3D Printing for Low-Cost and Versatile Attenuated Total Reflection Infrared Spectroscopy. Anal. Chem. 2020; 92, 7, 4736–4741. doi: 10.1021/acs.analchem.9b04043.

17. Baumgartner B, Freitag S, Gasser C, Lendl B. A pocket-sized 3D-printed attenuated total reflection-infrared filtometer combined with functionalized silica films for nitrate sensing in water. Sens. Actuators B Chem. 2020; 310, 127847. doi: 10.1016/j.snb.2020.127847.

18. Da Silva Junior JH, Castro PS. Enhancing analytical capabilities at an affordable cost: A lab-made 3D-printed external reflection accessory for versatile spectroscopic exploration. Spectrochim. Acta A Mol. Biomol. Spectrosc. 2025; 343, 15, 126585. doi: 10.1016/j.saa.2025.126585.

19. Bosman AJ, Freitag S, Ross GMS, Sulyok M, Krska R, Ruggeri FS, et al. Interconnectable 3D-printed sample processing modules for portable mycotoxin screening of intact wheat. Anal. Chim. Acta; 2024, 1285, 2, 342000. doi: 10.1016/j.aca.2023.342000.

20. Cruz-Tirado JP, Vieira MSS, Amigo JM, Siche R, Barbin DF. Prediction of protein and lipid content in black soldier fly (Hermetia illucens L.) larvae flour using portable NIR spectrometers and chemometrics. Food Control; 2023, 153, 109969. doi: 10.1016/j.foodcont.2023.109969.

21. Eady M, Payne M, Sortijas S, Bethea E, Jenkins D. A low-cost and portable near-infrared spectrometer using open-source multivariate data analysis software for rapid discriminatory quality assessment of medroxyprogesterone acetate injectables. Spectrochim Acta A Mol Biomol Spectrosc; 2021, 259, 119917. doi: 10.1016/j.saa.2021.119917.

22. Innospectra Corporation. NIR-M-R3® Reflective Module. [cited 07 December 2023]. In: Innospectra [internet]. Taiwan: Innospectra Corporation. Available from: https://www.inno-spectra.com/product_1_detail_a04#anchor_pt1_a04.

23. Burns DA, Ciurczak EW. Handbook of Near-Infrared Analysis. 3rd ed. Boca Raton: 966 CRC Press, Taylor & Francis Group, 2007.

24. Rego G, Ferrero F, Valledor M, Campo JC, Forcada S, Royo LJ, et al. A portable IoT NIR spectroscopic system to analyse the quality of dairy farm forage. Comput. Electron. Agric. 2020; 175, 105578, 1–8. doi: 10.1016/j.compag.2020.105578.

25. Varga E, Glauner T, Köppen R, Mayer K, Sulyok M, Schuhmacher R, et al. Stable isotope dilution assay for the accurate determination of mycotoxins in maize by UHPLC-MS/MS. Anal. Bioanal. Chem. 2012; 402, 2675–2686. doi: 10.1007/s00216-012-5757-5.

26. Stevens A, Ramirez-Lopez L & Viscarra Rossel RA (2022). prospectr: Miscellaneous functions for processing and sample selection of vis-NIR diffuse reflectance data (R package version 0.2.6) [Computer software]. https://CRAN.R-project.org/package=prospectr.

27. Wickham, H. (2023). ggplot2: Create elegant data visualizations using the grammar of graphics (R package version 3.5.1) [Computer software]. https://ggplot2.tidyverse.org.

28. Kassambara, A. (2023). ggpubr: ‘ggplot2’ based publication ready plots (R package version 0.6.0) [Computer software]. https://rpkgs.datanovia.com/ggpubr/.

29. Kucheryavskiy, S. (2023). mdatools: Multivariate data analysis for chemometrics (R package version 0.12.0) [Computer software]. https://github.com/svkucheryavski/mdatools.

30. Ciurczak EW, Igne B, Workman Jr., J., & Burns DA. (Eds.). (2021). Handbook of Near-Infrared Analysis. 4th ed. Boca Raton: 938 CRC Press, Taylor & Francis Group, 2021.

31. Zanotto D. (2022). Determination of geometric mean diameter of corn particle through near infrared spectroscopy (NIR). Proceedings of the 17th International Conference on Near Infrared Spectroscopy, Foz do Iguaçu, Brazil. doi: 10.1234/abcde.

32. Evans, CE, Frempong NS, Nortey TN, Stark CR, Chad BP (2021). The influence of ingredients, corn particle size, and sample preparation on the predictability of near infrared reflectance spectroscopy. Kansas Agricultural Experiment Station Research. Reports, 7(10). doi: 10.4148/2378-5977.8147.

33. Ezenarro J, Schorn-García D. How Are Chemometric Models Validated? A Systematic Review of Linear Regression Models for NIRS Data in Food Analysis. J. Chemom. 2025; 39, 6, e70036. doi: 10.1002/cem.70036.

34. Bailly S, Orlando B, Brustel J, Bailly, J-D., & Levasseur-Garcia C. Rapid detection of aflatoxins in ground maize using near infrared spectroscopy. Toxins. 2024; 16, 9, 1–12, Article 385. doi: 10.3390/toxins16090385

35. Yang Q, Li Y, Li J, Zhang Z, Liu Q, Guo G, et al. Optimizing Near-Infrared Spectroscopy Models for Rapid and Green Detection of Crude Protein and Fat in Crop Grains Using Sample Set Division. ACS Omega. 2025; 10, 15, 14755–14769. doi: 10.1021/acsomega.4c09155.

36. Liu Y, Liu Y, Chen Y, Zhang Y, Shi T, Wang J, et al. The Influence of Spectral Pretreatment on the Selection of Representative Calibration Samples for Soil Organic Matter Estimation Using Vis-NIR Reflectance Spectroscopy. Remote Sens. 2019; 11, 450. doi: 10.3390/rs11040450.

37. Lopez E, Etxebarria-Elezgarai J, Amigo JM, Seifert A. The importance of choosing a proper validation strategy in predictive models. A tutorial with real examples. Anal. Chim. Acta. 2023; 1275, 22, 341532. doi: 10.1016/j.aca.2023.341532.

38. Van den Goorbergh R, Van Smeden M, Timmerman D, Van Calster B. The harm of class imbalance corrections for risk prediction models: illustration and simulation using logistic regression. J Am Med Inform Assoc. 2022; 16, 29(9), 1525-1534. doi: 10.1093/jamia/ocac093.

39. Barker M, Rayens W. Partial least squares for discrimination. J. Chemom. 2003; 17, 3, 166– 173. doi: 10.1002/cem.785.

40. Freitag S, Anlanger M, Fomina P, Femenias A, Aledda M, Mizaikoff B, et al. Attenuated total reflection mid-infrared spectroscopy to screen Austrian and French wheat from multiple years for deoxynivalenol. Spectrochim. Acta A Mol. Biomol. Spectrosc. 2025; 340, 5, 126340. doi: 10.1016/j.saa.2025.126340.

41. Von Holst C, Lattanzio VMT (2025). Rapid mycotoxin screening: An update on the evaluation of analytical performances and relevant criteria in relation to the method scope. In: Suman M (edit.). Mycotoxins: Screening and confirmatory analysis (p. 4–170). Brill (Wageningen Academic), 2. doi: 10.1163/9789004724969_003.

42. Mansuri SM, Chakraborty S, Pandiselvam R, & colab. Effect of germ orientation during Vis–NIR hyperspectral imaging for the detection of fungal contamination in maize kernel using PLS-DA, ANN and 1D-CNN modelling. Food Control. 2022; 139, 109077. doi: 10.1016/j.foodcont.2022.109077.

43. Borràs-Vallverdú B, Marín S, Sanchis, V, Gatius, F, & Ramos AJ. NIR-HSI as a tool to predict deoxynivalenol and fumonisins in maize kernels: a step forward in preventing mycotoxin contamination. J. Sci. Food Agric. 2024; 104, 9, 5495–5503. doi: 10.1002/jsfa.13388.

44. Parrag V, Gillay Z. Hyperspectral imaging application to determine mycotoxin content in cornmeal samples. Prog. Agric. Eng. Sci. 2020; 16, 1, 51–60. doi: 10.1556/446.2020.00009.

45. Liang K, Liu QX, Xu JH, Wang YQ, Okinda CS, Shena MX. Determination and Visualization of Different Levels of Deoxynivalenol in Bulk Wheat Kernels by Hyperspectral Imaging. J Appl Spectrosc. 2018; 85, 953–961. doi: 10.1007/s10812-018-0745-y.

46. Liang K, Huang J, He R, Wang Q, Chai Y, Shen M. Comparison of Vis-NIR and SWIR hyperspectral imaging for the non-destructive detection of DON levels in Fusarium head blight wheat kernels and wheat flour. Infrared Phys. Technol. 2020; 106, 103281. doi: 10.1016/j.infrared.2020.103281.

47. Palumbo R, Crisci A, Venâncio A, Cortiñas Abrahantes J, Dorne JL, Battilani P, et al. Occurrence and Co-Occurrence of Mycotoxins in Cereal-Based Feed and Food. Microorganisms. 2020; 8, 1, 74. doi: 10.3390/microorganisms8010074.

48. Femenias A, Bainotti MB, Gatius F, Ramos AJ, & Marín S. Standardization of near infrared hyperspectral imaging for wheat single kernel sorting according to deoxynivalenol level. Food Res Int. 2021; 139, 109925. doi: 10.1016/j.foodres.2020.109925.

49. Tyska D, Mallmann AO, Vidal JK, Almeida CAA de, Gressler LT, Mallmann CA. Multivariate method for prediction of fumonisins B1 and B2 and zearalenone in Brazilian maize using Near Infrared Spectroscopy (NIR). PLoS ONE. 2021; 16, 1, 0244957, 1-14. doi: 10.1371/journal.

50. Caramês ETS, Piacentini KC, Alves LT, Pallone JAL, Rocha LO. NIR spectroscopy and chemometric tools to identify high content of deoxynivalenol in barley. Food Addit. Contam. Part A. 2020; 37, 9, 1542–1552. doi: 10.1080/19440049.2020.1778189.

51. Lim J, Kim G, Mo C, Oh K, Yoo H, Ham H. et al. Classification of fusarium-infected korean hulled barley using near-infrared reflectance spectroscopy and partial least squares discriminant analysis. Sensors. 2017; 17, 2258. doi:10.3390/s17102258.

52. Falade TDO, Sultanbawa Y, Fletcher MT, & Fox G. Near Infrared Spectrometry for Rapid Non-Invasive Modelling of Aspergillus-Contaminated Maturing Kernels of Maize (Zea mays L.). Agriculture. 2017; 7, 9, 77. doi: 10.3390/agriculture7090077.

53. Vicens-Sans A, Pascari X, Molino F, Ramos AJ, Marín S. Near infrared hyperspectral imaging as a sorting tool for deoxynivalenol reduction in wheat batches. Food Res. Int. 2024; 178, 113984. doi: 10.1016/j.foodres.2024.113984.

54. Ellison, SLR, Williams A (Eds.). (2012). Quantifying uncertainty in analytical measurement (3rd ed.). EURACHEM/CITAC. Avaiable in: https://www.eurachem.org/images/stories/Guides/pdf/QUAM2012_P1.pdf.

55. Pascale M, De Girolamo A, Lippolis V, Stroka J, Mol HGJ, Lattanzio VMT. Performance Evaluation of LC-MS Methods for Multimycotoxin Determination. J. AOAC Int. 2019; 102, 6, 1708–1720. doi: 10.1093/jaoac/102.6.1708.

56. Whitaker TB. Sampling Foods for Mycotoxins. Food Addit. Contam. 2007; 23, 1, 50–61. doi: 10.1080/02652030500241587.

